# Determining Allele-Specific Protein Expression (ASPE) Using a QconCAT-Based Proteomics Method: a Novel Approach to Identify *Cis*-acting Genetic Variants

**DOI:** 10.1101/264614

**Authors:** Jian Shi, Xinwen Wang, Huaijun Zhu, Hui Jiang, Danxin Wang, Alexey Nesvizhskii, Hao-Jie Zhu

## Abstract

Measuring allele-specific expression (ASE) is a powerful approach for identifying *cis-*regulatory genetic variants. Here we developed a novel targeted proteomics method for quantification of allele-specific protein expression (ASPE) based on scheduled high resolution multiple reaction monitoring (sMRM-HR) with a heavy stable isotope-labeled quantitative concatamer (QconCAT) internal protein standard. This strategy was applied to the determination of the ASPE of UGT2B15 in human livers using the common *UGT2B15* nonsynonymous variant rs1902023 (i.e. Y85D) as the marker to differentiate expressions from the two alleles. The QconCAT standard contains both the wild type tryptic peptide and the Y85D mutant peptide at a ratio of 1:1 to ensure accurate measurement of the ASPE of UGT2B15. The results from 18 UGT2B15 Y85D heterozygotes revealed that the ratios between wild type Y allele and mutant D allele varied from 0.60 to 1.46, indicating the presence of *cis*-regulatory variants. In addition, we observed no significant correlations between the ASPE and mRNA ASE of *UGT2B15*, suggesting the involvement of different *cis*-acting variants in regulating the transcription and translation processes of the gene. This novel ASPE approach provides a powerful tool for capturing *cis*-genetic variants involved in post-transcription processes, an important yet understudied area of research.

## INTRODUCTION

Gene expression can be regulated by both *cis*- and *trans*-regulatory factors. *Cis*-regulatory genetic variants are nearby or within a gene locus that affect the gene expression in an allele-specific manner. The most well-established *cis*-acting variants are located in the enhancer and promoter regions of a gene ^1^. *Trans*-regulatory factors regulate gene expression of both alleles. They can be genetic variants, such as those of transcription factors genes, or non-genetic factors, such as drug inducers, disease states, and different developmental stages. Although still an area of debate, some recently published large-scale genomic studies suggest that *cis*-acting variants have greater impacts on gene expression variability than *trans*-acting variants ^2^. Because *cis*-regulatory factors have allele-specific effects, measuring allele-specific expression (ASE) has been widely used as a powerful approach to search for *cis*-acting variants. Deviation of an allelic expression ratio from one would strongly suggest the presence of *cis*-regulatory variant(s). Allelic expression analysis takes advantage of the fact that one allele serves as the control for the other, cancelling out the effects from *trans*-regulatory elements and non-genetic confounding factors. Thus, relative to total gene expression, ASE is unlikely to be altered by non-genetic factors and *trans*-regulatory elements, and thus is more sensitive and robust for the identification of functional *cis*-regulatory genetic variants.

Most allele-specific expression studies are conducted at the mRNA level due to technologies for the quantitative analysis of mRNA expressions being more amenable than those for protein expression measurement. However, mRNA expressions correlate poorly with protein expressions for many genes as protein abundance is subject to post-transcriptional regulations ^3–6^. Thus, regulatory variants identified through an mRNA ASE assay may not be predictive of protein expression in genes with poor mRNA-protein expression correlations. Furthermore, mRNA ASE analysis is unable to detect variants that affect gene expression at the post-transcriptional level (e.g. translation efficiency and protein degradation). Therefore, studies based on mRNA ASE cannot fully capture and predict the effects of *cis*-acting genetic variants on gene expression at the protein level.

In the present study, we developed a novel targeted proteomics assay for precise quantification of allele-specific protein expression (ASPE) using a scheduled high resolution multiple reaction monitoring (sMRM-HR) approach with a heavy stable isotope-labeled quantitative concatamer (QconCAT) internal protein standard. The UGT2B15 gene, which encodes the important phase II metabolizing enzyme UDP glucuronosyltransferase 2B15 expressed in human livers, was targeted in this proof-of-concept study. The mutant tryptic peptide N<u>D</u>LEDSLLK harboring the common UGT2B15 nonsynonymous variant rs1902023 (i.e. Y85D) and its wild type counterpart N<u>Y</u>LEDSLLK were quantified simultaneously to determine the ASPE ratios of the gene (Figure 1). This assay exhibited excellent precision and reproducibility, and has the potential to be widely utilized for the identification *cis*-regulatory genetic variants.

**Figure 1:**
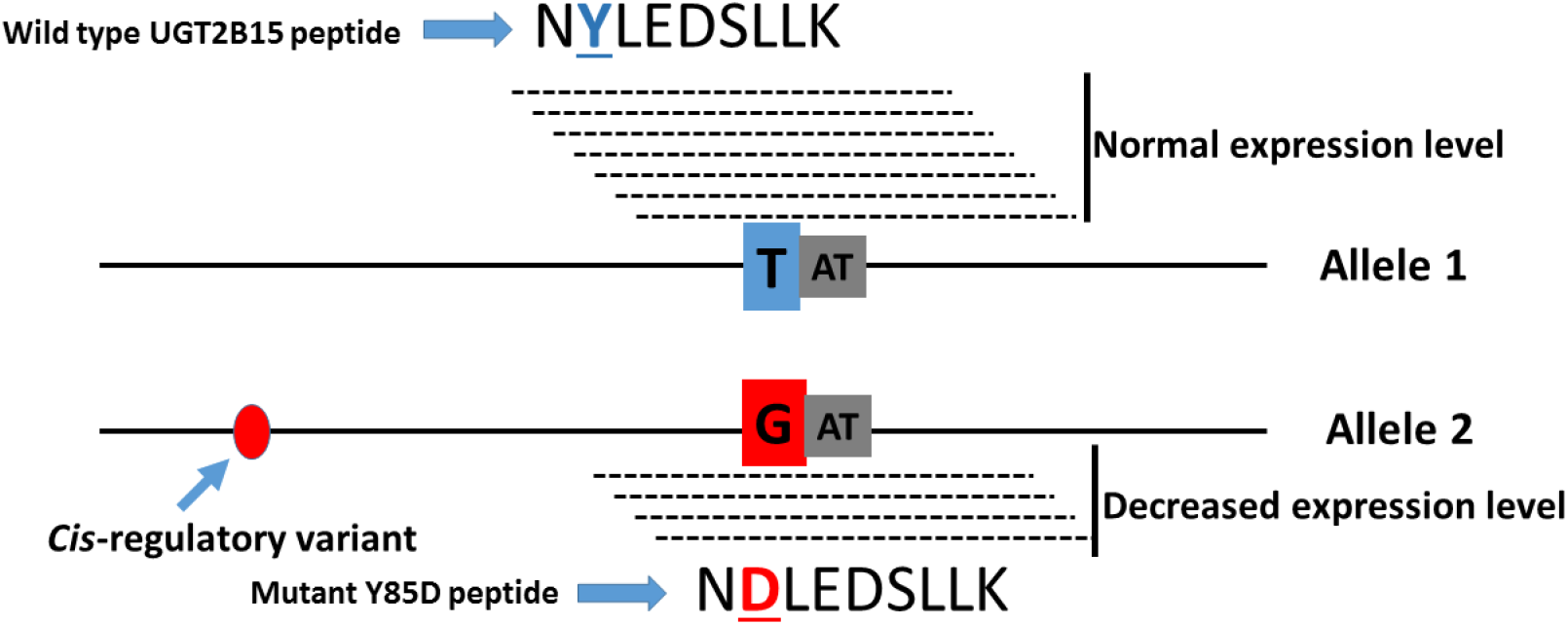
Schematic illustration of the ASPE approach for the detection of *cis*-regulatory variant of UGT2B15. Both the wild type tryptic peptide N<u>Y</u>LEDSLLK and the corresponding Y85D mutant peptide N<u>D</u>LEDSLLK are simultaneously quantified; and a ratio of Y-to-D ASPE deviating from one would suggest the existence of a *cis*-regulatory variant.

## MATERIALS AND METHODS

### Materials

Trifluoroacetic acid, formic acid, acetonitrile, isopropyl-D-1-thiogalactopyranoside disodium phosphate, M9 salts, magnesium sulfate, calcium chloride hexahydrate, sodium chloride, thiamine, glucose, imidazole, amino acids, and Benzonase^®^ nuclease were purchased from Sigma-Aldrich (St. Louis, MO). ^13^C6 arginine and ^13^C6, ^15^N2 lysine were obtained from Cambridge Isotope Laboratories (Tewksbury, MA). Urea and dithiothreitol were purchased from Fisher Scientific Co. (Pittsburgh, PA). Iodoacetamide and ammonium bicarbonate were the products of Acros Organics (Morris Plains, NJ). TPCK-treated trypsin was obtained from Worthington Biochemical Corporation (Freehold, NJ). Lysyl endopeptidase was purchased from Wako Chemicals (Richmond, VA). Water Oasis HLB columns were from Waters Corporation (Milford, MA). Synthetic iRT standards solution was from Biognosys AG (Cambridge, MA). Escherichia coli strain BL21(DE3) and BugBuster^®^ protein extraction reagent were products of EMD Millipore (Burlington, MA). HisTrap HP histidine-tagged protein purification columns were from GE Healthcare (Pittsburgh, PA). Lysozyme Solution (50 mg/mL), slide-A-Lyzer G2 dialysis cassettes (3.5K MWCO) and PierceTM BCA protein assay kit were obtained from ThermoFisher Scientific (Waltham, MA). Normal human liver samples were obtained from XenoTech LLC (Kansas City, KS) and the Cooperative Human Tissue Network (Columbus, OH). All other chemicals and reagents were of analytical grade and commercially available.

### UGT2B15 genotyping

A total of 40 individual human liver samples were genotyped for the *UGT2B15* nonsynonymous variant rs1902023 (Y85D) based on a previously published pyrosequencing protocol with some modifications ^7^. Briefly, genomic DNA was extracted from human liver tissues using the PureLink^®^ Genomic DNA Mini Kit (Thermo Fisher Scientific Co., Fair Lawn, NJ) according to the manufacturer’s instructions. Polymerase chain reaction (PCR) was carried out to amplify the target region of the variant using the primer pair 5’-TCAATGCCAGTAAATCATCTGC-3’ (biotinylated) and 5’-TCGAGAATTTTCAGAAGAGAATCT-3’. The biotinylated strand of the amplified DNA was subjected to prosequencing using the sequencing primer 5’-TCAGAAGAGAATCTTCCAAA-3’ on a PyroMark Q96 MD instrument (Qiagen, Valencia, CA) following a standard protocol. Genotyping data were analyzed using the PyroMark Q96 MD software.

### UGT2B15 QconCAT protein standard design and generation

The unique UGT2B15 peptide N<u>Y</u>LEDSLLK and its variant peptide N<u>D</u>LEDSLLK (i.e., Y85D, rs1902023) were chosen as the markers for the quantification of ASPE of UGT2B15 with a QconCAT internal standard in human livers. It is important that the internal standard includes equimolar amounts of the wild type and mutant QconCAT peptides in order to precisely quantify the ASPE of a gene. Accordingly, we designed a QconCAT protein containing both N<u>Y</u>LEDSLLK and N<u>D</u>LEDSLLK peptides at a ratio of 1:1. In addition, both peptides are flanked by at least 15 native UGT2B15 amino acids, such that trypsin digestion will result in identical digestion efficiencies for the heavy QconCAT peptides and unlabeled endogenous peptides from human livers. The amino acid sequences and features of the QconCAT protein are shown in Figure 2.

**Figure 2:**
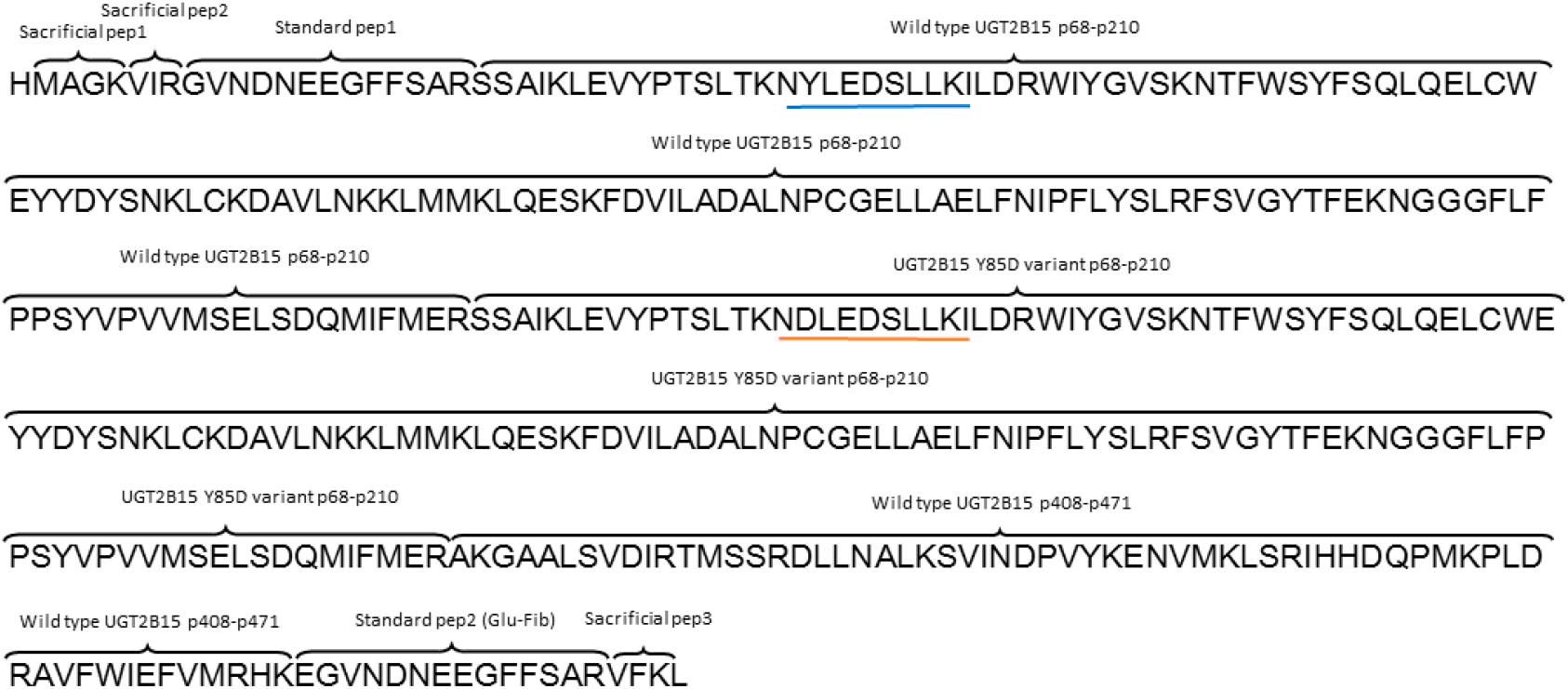
Amino acid sequences and features of the UGT2B15 QconCAT standard. The wild type tryptic peptide N<u>Y</u>LEDSLLK and the corresponding Y85D mutant peptide N<u>D</u>LEDSLLK are highlighted with blue and orange underlines, respectively. Both tryptic peptides are flanked by at least 15 native UGT2B15 amino acids to enable identical digestion efficiencies for both the heavy QconCAT peptides and the light peptides from human liver samples. Other features of the QconCAT standard include three sacrificial peptides and two standard peptides for the quantification of the QconCAT concentrations.

The QconCAT DNA construct was synthesized de novo and cloned into the pET21b plasmid by Synbio Technologies (Monmouth Junction, NJ). Escherichia coli strain BL21(DE3) from Novagen was transformed with the plasmid and cultured in minimal medium (M9 salts, 1 mmol/L MgSO_4_, 0.1 mmol/L CaCl_2_, 0.00005% [w/v] thiamine, and 0.2% [w/v] glucose) supplemented with ^13^C_6_ arginine, ^13^C_6_, ^15^N_2_ lysine (0.2 mg/mL, Cambridge Isotope Laboratories, Tewksbury, MA), proline, histidine, tyrosine, phenylalanine, tryptophan (0.2 mg/mL, Sigma-Aldrich), and the remaining 13 amino acids (0.1 mg/mL, Sigma-Aldrich). When cells were grown to mid log phase (A_600_ 0.6-0.8), QconCAT protein expression was induced by adding 1 mmol/L isopropyl-D-1-thiogalactopyranoside. After 5 hours of growth at 37 °C, cells were harvested by centrifugation and processed using a reported method with some minor modifications ^8^. Briefly, cells were lysed with BugBuster^®^ protein extraction reagent (EMD Millipore) and Benzonase^®^ nuclease (Sigma-Aldrich). The resulted inclusion bodies were suspended in 20 mmol/L disodium phosphate, 8 mol/L urea, 0.5 mol/L NaCl, 20 mmol/L imidazole (pH 7.4). QconCAT proteins were then purified using affinity chromatography with HisTrap HP columns (GE Healthcare Bio-Sciences, Pittsburgh, PA). The purified QconCAT proteins were desalted after three rounds of dialysis using Slide-A-Lyzer G2 dialysis cassettes (3.5K MWCO, Thermo Fisher Scientific, Waltham, MA) against 50 mmol/L ammonium bicarbonate containing 1 mmol/L dithiothreitol. The QconCAT standards were stored at −80 °C until use.

### Scheduled high resolution multiple reaction monitoring (sMRM-HR) analysis of UGT2B15 ASPE in human liver microsomes

To prepare human liver microsomes, approximate 200 mg frozen liver tissues were homogenized in 0.6 mL ice-cold phosphate-buffered saline (PBS) in 1.5 mL microcentrifuge tubes using a microtube homogenizer (VWR International LLC, Chicago, USA). S9 fractions were obtained after centrifugation of the homogenates at 10,000 g for 30 min at 4 °C. The supernatants (S9 fractions) were transferred to Beckman ultracentrifuge tubes and centrifuged at 300,000 g (80,000 rpm) for 20 min. Microsomes were obtained by resuspending the pellets in PBS using a microtube homogenizer. Protein concentrations of the microsomes were determined using a Pierce™ BCA protein assay kit (Thermo Fisher Scientific, Waltham, MA). The microsome samples were stored at −80 °C until use.

A previous reported Lys-C/Trypsin combinatorial digestion protocol was utilized for protein digestion, with some modifications ^9^. An aliquot of 80 μg microsome protein was mixed with a 10-fold volume of pre-cooled acetone. The mixture was briefly vortexed and incubated at −20 °C for at least 2 hours, followed by centrifugation at 17,000 *g* for 15 min at 4 °C. After removing the supernatants, the precipitated proteins were washed with 200 μL ice-cold 80% ethanol, and centrifuged again at 17,000 *g* for 15 min at 4 °C. The supernatants were discarded, and the precipitated proteins were air-dried at room temperature before being resuspended in 92.5 μL of freshly prepared 4 mM dithiothreitol in 8 M urea solution containing 100 mM ammonium bicarbonate. Next, the heavy stable isotope-labeled QconCAT UGT2B15 internal standard (75 ng) was added. Samples were then briefly vortexed and sonicated, and incubated at 37 °C for 45 min. After samples were cooled down to room temperature, 100 μL of freshly prepared 20 mM iodoacetamide in 8 M urea/100 mM ammonium bicarbonate solution was added. The mixture was incubated at room temperature for 30 min in the dark for alkylation.

Following incubation, the urea concentration was adjusted to 6 M by adding 64.6 μL of 50 mM ammonium bicarbonate. Lysyl endopeptidase (Wako Chemicals, Richmond, VA) was then added for the first step digestion at 37 °C for 6 h (protein to lysyl endopeptidase ratio = 100:1). Samples were diluted with 50 mM ammonium bicarbonate to further decrease urea concentration to 1.6 M, and then subjected to the second step of digestion with trypsin overnight at 37 °C at a protein to trypsin ratio of 50:1. Digestion was terminated by the addition of 1 μL trifluoroacetic acid. Digested peptides were extracted and purified using Waters Oasis HLB columns according to the manufacturer’s instructions. Peptides eluted from the columns were dried in a SpeedVac SPD1010 concentrator (Thermo Scientific, Hudson, NH), and reconstituted in 80 μL of 3% acetonitrile solution containing 0.1% formic acid. The peptide samples were centrifuged at 17,000 *g* for 10 min at 4 °C, and 40 μL of the supernatant was collected and supplemented with 1 μL of the synthetic iRT standards solution (Biognosys AG, Cambridge, MA) for LC-MS/MS analysis.

Information-dependent acquisition (IDA) was performed on three pooled human liver microsome samples using the method described in a recently published study ^10^ to generate the reference spectral library for the sMRM-HR data analysis. A sMRM-HR analysis was performed on a TripleTOF 5600+ mass spectrometer (AB Sciex, Framingham, MA) for the quantification of both the wild type peptide N<u>Y</u>LEDSLLK and the mutant peptide N<u>D</u>LEDSLLK. An Eksigent 2D plus LC system (Eksigent Technologies, Dublin, CA) was utilized for peptide separation using a trap-elute configuration, which included a trapping column (ChromXP C18-CL, 120 Å, 5 μm, 0.3 mm cartridge, Eksigent Technologies, Dublin, CA) and an analytical column (ChromXP C18-CL, 120 Å, 150 × 0.3 mm, 5 μm, Eksigent Technologies, Dublin, CA). The mobile phase consisted of water with 0.1% formic acid (phase A) and acetonitrile containing 0.1% formic acid (phase B). Peptides were trapped and cleaned on the trapping column with the mobile phase A delivered at a flow rate of 10 μL/min for 3 min followed by the separation on the analytical column with a 32 min gradient at a flow rate of 5 μL/min. The gradient time program was set as follows for the phase B: 0 to 22 min: 3% to 30%, 22 to 25 min: 30% to 40%, 25 to 26 min: 40% to 80%, 26 to 27 min: 80%, 27 to 28 min: 80% to 3%, and 28 to 32 min at 3% for column equilibration.

The sMRM-HR acquisition consisted of one 200 ms TOF-MS scan from 400 to 1250 Da and subsequent MS/MS scans from 100 to 1500 Da (50 ms accumulation time, 50 mDa mass tolerance, charge state from +2 to +5 which exceeds 2,000,000 cps, 30 maximum candidate ions to monitor per cycle, rolling collision energy) of the inclusion precursors with the scheduled retention times. The retention times were obtained from pilot non-scheduled MRM-HR runs. The intensity threshold of the target precursors was set to 0 and scheduling window was defined as 270 s. The target peptides/precursors and their retention times are summarized in Supplemental Table 1.

The sMRM-HR data were analyzed using the Skyline software (version 3.7.1.11271, University of Washington, Seattle, WA) ^11^ with automatic detections of MS/MS chromatographic peaks against the spectral library generated from the data obtained inthe Information Dependent Acquisition mode. The selected peaks were reviewed manually after the automated analysis. Both MS1 and MS/MS filtering were set as a “TOF” mass analyzer with the resolution power of 30,000 and 15,000, respectively. Peak areas of the top four fragment ions were summed up and normalized to the isotope-labeled QconCAT internal standards for peptide quantification. All LC-MS/MS data have been deposited to the ProteomeXchange Consortium via the PRIDE ^12^ partner repository with the dataset identifier PXD008788.

### UGT2B15 mRNA allele-specific expression analysis using SNapShot

Samples heterozygous for rs1902023 were also analyzed for allele-specific expression at the mRNA level using a SNaPshot method ^13^. Briefly, *UGT2B15* cDNA was obtained from reverse-transcription, and a fragment of DNA and *UGT2B15* cDNA surrounding rs1902023 was amplified using PCR. PCR primers are listed in Supplemental Table 2. After degradation of un-incorporated dNTPs and excess primers with exonuclease I and Antarctic alkaline phosphatase, PCR products were then subjected to a primer extension assay (SNaPshot kit, Life Tech, Grand Island, NY) using extension primers designed to anneal to the amplified DNA adjacent to the rs1902023 site. The resulting primer extension products were then analyzed on an ABI3730 capillary electrophoresis DNA instrument using Gene Mapper software (Life Tech, Grand Island, NY). Two independent measurements, each in duplicates, were performed.

## RESULTS

Out of the 40 liver samples genotyped by Pyrosequencing, a total of 18 were found to be heterozygous for the *UG2B15* variant Y85D. A heavy isotope-labeled standard was successfully generated using a QconCAT approach (Supplemental Figure 1), which contains both the wild type peptide N<u>Y</u>LEDSLLK and the mutant counterpart N<u>D</u>LEDSLLK harboring the Y85D variant. The QconCAT standard was applied in a sMRM-HR study, and the expression levels of two alleles of *UGT2B15* were determined based on the ratios of the light peptides N<u>Y</u>LEDSLLK and N<u>D</u>LEDSLLK to the corresponding heavy internal standards. UGT2B15 ASPE was quantified according to the expression ratios of the wild type peptide to the mutant peptide of the Y85D. The sMRM-HR method with heavy isotope-labeled internal standards exhibited excellent sensitivity and selectivity for the quantification of both peptides, as shown in the chromatograms in Supplemental Figure 2. The ASPE ratios of the 18 samples ranged from 0.61 to 1.42 with an average of 1.05 (Figure 3). To evaluate the reproducibility of the assay, the first six samples were independently measured twice. The measurements were found to be highly reproducible with relative standard deviations less than 8% for all samples (Figure 4). The mRNA ASE of *UGT2B15* were quantified using a SNapShot method based on the same genetic marker Y85D (rs1902023). The mRNA ASE ratios were between 0.68 and 1.24 with a mean value of 0.89. However, no significant correlation was found between the ASPE and the mRNA ASE ratios (Figure 5).

**Figure 3:**
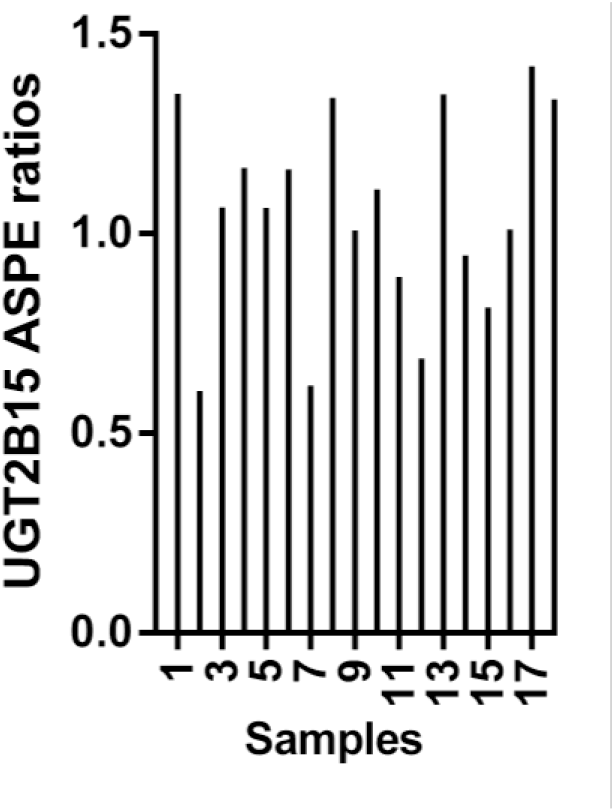
Distribution of the ASPE ratios of UGT2B15 determined in 18 human liver samples with the heterozygous Y85D genotype. The abundances of the wild type peptide N<u>Y</u>LEDSLLK (Y allele) and the mutant peptide N<u>D</u>LEDSLLK (D allele) were calculated based on the ratios of the light peptides to the heavy QconCAT internal standard peptides. The Y to D ASPE ratios ranged from 0.61 to 1.42 among the 18 Y85D heterozygous samples.

**Figure 4.**
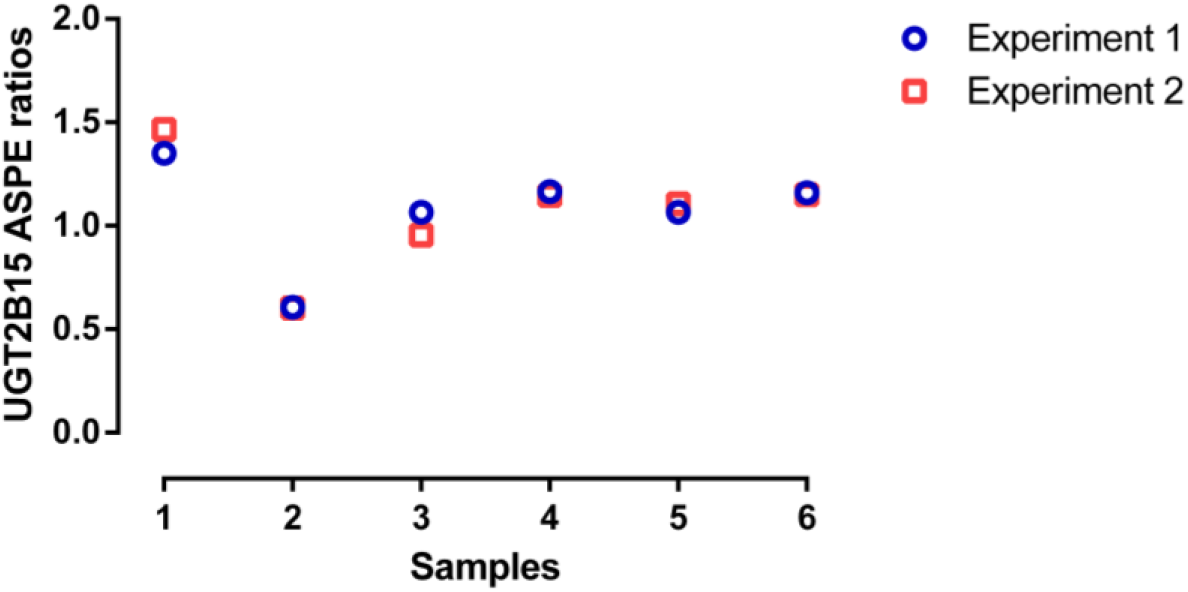
Reproducibility of the novel QconCAT-based sMRM-HR proteomics assay for the measurement of UGT2B15 ASPE ratios. The data are the measurements from two independent experiments of six human liver samples. The relative standard deviations were less than 8% for all samples.

**Figure 5.**
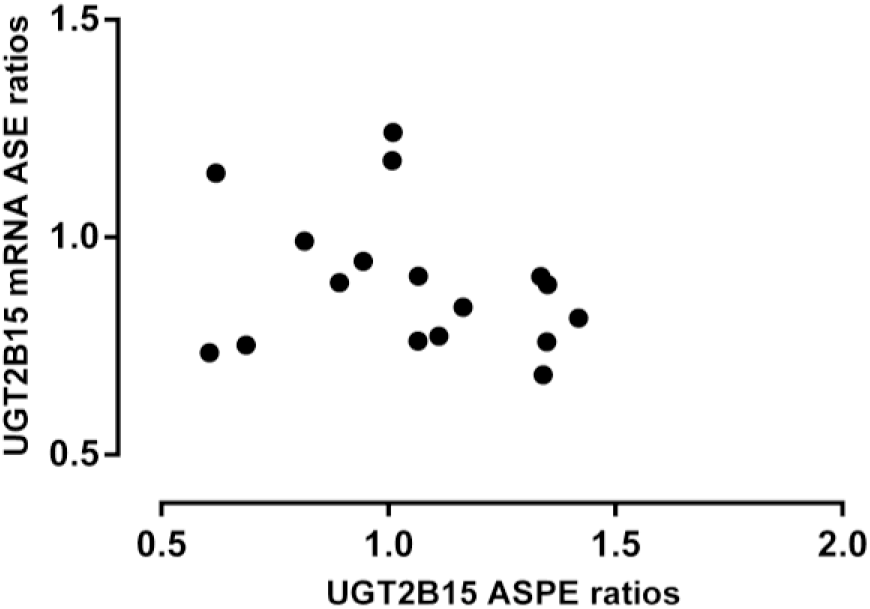
No significant correlation between the ASPE and mRNA ASE ratios of UGT2B15 in human livers. The ASPE and mRNA ASE of UGT2B15 were quantified using the QconCAT-based sMRM-HR proteomics assay and a SNapShot assay, respectively, in 18 Y85D heterozygotes. The ASPE ratios do not significantly correlate with the ratios of mRNA ASE (p = 0.27).

## DISCUSSION

We have developed a QconCAT-based targeted proteomics approach for the measurement of ASPE with high precision and reproducibility. Applying this approach to determining UGT2B15 ASPE in human livers, we revealed a robust allelic difference in protein expression of UGT2B15. The ASPE cannot be explained by mRNA ASE, indicating the presence of *cis*-acting regulatory variants affecting UGT2B15 expression at the post-transcriptional level. The results demonstrate that this novel ASPE approach has the potential to be widely utilized for the identification of *cis*-acting genetic variants that can be difficult to be detected by conventional mRNA and/or protein expression-based methods.

Despite extensive research in the past several decades, a significant portion of inheritable phenotypes for many genes cannot be explained by the functional genetic variants identified so far. For example, the catalytic activity of CYP3A4, one of the most abundantly expressed phase I enzymes in human livers, is highly correlated to protein expression and markedly inheritable (66% − 88%) according to both an early twin study and a more recent study using a repeated drug administration approach ^14, 15^. However, all identified functional genetic variants can only explain less than 30% of the variability in CYP3A4 function. The gap between the expected and known genetic contributions to phenotypic traits is termed “missing heritability” ^16^. Identifying genetic variants that accounts for “missing heritability” is paramount for the understanding of interindividual variability in various phenotypes, such as diseases susceptibility and response to drug therapy.

The most commonly used approach to identify regulatory genetic variants is to measure total mRNA expression or mRNA ASE and then search for genetic variants associated with the mRNA expression or ASE imbalance. However, numerous studies have demonstrated that the correlations between mRNA and protein expressions are very poor for many genes ^3–6^. Genetic variants associated with altered mRNA expression levels may not necessarily affect protein expression of the gene (i.e. false positive) ^3^, which could be partially due to a stronger evolutionary constraint at protein level relative to mRNA expression ^17^. Furthermore, genetic polymorphisms affecting gene expression at the protein level may not be readily detected by mRNA expression-based approaches (i.e. false negative) ^4, 18, 19^. For example, several linked variants were strongly associated with protein levels of the apolipoprotein L. 2 (*APOL2*) gene, but the effect was unrelated to either mRNA or ribosome occupancy ^4^. It is not uncommon that functional genetic variants identified from an mRNA expression study were later found to have no observable effects on protein expression and/or clinical phenotypes. The lack of correlation between mRNA and protein expressions may in part explain this discrepancy.

In recent years, with the progress of mass spectrometry-based quantitative proteomics, protein expression has been increasingly utilized as a phenotype for the study of functional regulatory genetic variants ^4, 20^. In most biological processes, protein instead of mRNA is the functional molecule; thus genetic variants associated with protein expression levels are more likely to be biologically relevant. It should be noted that in addition to genetic variants, many non-genetic factors such as inducers and disease state can also affect protein expression ^21^. Therefore, observation of elevated or decreased protein expression doesn’t necessarily suggest the existence of functional regulatory genetic variants. Consequently, using protein expression level as the phenotype for identifying functional genetic variants can be significantly confounded by non-genetic regulatory elements, which may significantly impair the statistical power of that approach and lead to false positive or negative findings.

Both *cis*- and *trans*-acting elements contribute to interindividual variability in gene expression. *Cis*-acting variants affect gene expression in an allele-specific manner, resulting in an ASPE ratio deviating from one. In contrast, *trans*-regulatory variants and environmental factors affect gene expression of both alleles, thus the ASPE ratio remains one. Therefore, measuring ASPE could be a powerful approach for the study of *cis*-regulatory variants because it is insusceptible to the confounding effects of *trans*-elements and environmental contributors. Many *cis*-elements, such as those in the promoter and enhancer regions, can lead to imbalanced ASPE through affecting mRNA allelic expression. In addition, some *cis*-regulatory variants can regulate gene expression at the post-transcriptional level, which include the variants causing changes to upstream open reading frame and the sequences around the translation initiation site ^18^ i addition to the variants associated with protein stability ^22^. In principle, this type of variants cannot be identified by mRNA ASE-based studies, but will be readily detected by an ASPE assay. ASPE has been quantified in several proteomics studies, which revealed wide spread contributions of *cis*-regulatory factors to variability in protein expression ^4, 23, 24^. However, the data-dependent acquisition (DDA) or label-free proteomics techniques utilized in these studies exhibit intrinsic limitations in the accuracy, precision, and repeatability of protein quantification ^25^. In the present study, we developed a novel sMRM-HR method using a heavy stable isotope-labeled QconCAT protein standard for the measurement of UGT2B15 ASPE in human liver. *UGT2B15* was chosen as the model gene for this proof-of-concept ASPE study due to 1) playing an important role in the metabolism of some endobiotics, such as steroid hormones, and several clinically important medications; 2) its protein expression varies markedly among individuals ^5^, however, genetic variants responsible for the expression variability remain largely undetermined; and 3) the nonsynonymous variant Y85D (rs1902023) is very common, with a minor allele frequency ranging from 44% to 64% in different populations, allowing adequate numbers of Y85D heterozygotes to be obtained from our banked human liver samples.

Combined with the use of heavy isotope-labeled internal standards, targeted proteomics approaches, such as the sMRM-HR method used in the present study enable the most accurate and reproducible protein quantification relative to other untargeted and/or unlabeled proteomics methods. QconCAT-based targeted proteomics haves been used for the quantification of many proteins including UGT2B15 ^26, 27^. The QconCAT protein standard we designed differs from conventional QconCAT standard in two important aspects. First, the standard contains both the wild type surrogate peptide N<u>Y</u>LEDSLLK and the mutant Y85D peptide N<u>D</u>LEDSLLK at a ratio of 1:1, which ensures that the heavy internal standard contains equimolar amounts of the two peptides. Second, both the wild type and mutant peptides are flanked by at least 15 native UGT2B15 amino acids to enable identical digestion efficiencies for both the heavy QconCAT peptides and the light peptides from human liver samples (Figure 2). The digestion enzyme trypsin interacts with three to four amino acid residues around scissile bonds, and certain residues, such as negatively charged amino acids, can significantly affect the digestion efficiency of trypsin ^28, 29^. Thus, digestion efficiency of the concatenated surrogate peptides in a conventional QconCAT protein can differ significantly from that of the peptides in native proteins, which imposes significant challenges on the accuracy and repeatability of QconCAT-based quantitative proteomics. Cheung et al. assessed the effect of natural amino acid flanking sequence on trypsin digestion efficiency of QconCAT proteins and concluded that including six or more amino acid flanking residues is necessary for reliable quantification ^30^. In the preset study, we included at least 15 native flanking amino acids to ensure accurate measurement of any departure of ASPE ratios from one. This new approach has demonstrated excellent sensitivity and reproducibility, and revealed a significant ASPE imbalance of UGT2B15 in human livers, with the Y-to-D allele ratios from 0.61 to 1.42, indicating the presence of *cis*-regulatory variant(s) of the gene. In addition, no significant correlation was observed between the ASPE and mRNA ASE, suggesting the existence of post-transcriptional regulatory variants capable of regulating UGT2B15 protein expression independent of the variants causing the mRNA ASE imbalance. This observation is consistent with a previous study showing that *UGT2B15* mRNA expressions did not significantly correlate to its protein levels in human livers ^5^. These results highlight the utility of this novel ASPE assay in determining genetic variants involved in post-transcriptional regulation, and further emphasize the importance of studying gene expression regulation at the protein level.

## Acknowledgements

This work was partially supported by the University of Michigan MCubed program and the National Institutes of Health National Heart, Lung, and Blood Institute [Grant R01HL126969, Hao-Jie Zhu].

## References

1. Wittkopp, P. J.; Kalay, G., Cis-regulatory elements: molecular mechanisms and evolutionary processes underlying divergence. Nature reviews. Genetics 2012, 13, (1), 59–69.

2. Osada, N.; Miyagi, R.; Takahashi, A., Cis- and *Trans*-regulatory Effects on Gene Expression in a Natural Population of Drosophila melanogaster. Genetics 2017, 206, (4), 2139–2148.

3. Sanford, J. C.; Wang, X.; Shi, J.; Barrie, E. S.; Wang, D.; Zhu, H. J.; Sadee, W., Regulatory effects of genomic translocations at the human carboxylesterase-1 (CES1) gene locus. Pharmacogenet Genomics 2016, 26, (5), 197–207.

4. Battle, A.; Khan, Z.; Wang, S. H.; Mitrano, A.; Ford, M. J.; Pritchard, J. K.; Gilad, Y., Genomic variation. Impact of regulatory variation from RNA to protein. Science 2015, 347, (6222), 664–7.

5. Ohtsuki, S.; Schaefer, O.; Kawakami, H.; Inoue, T.; Liehner, S.; Saito, A.; Ishiguro, N.; Kishimoto, W.; Ludwig-Schwellinger, E.; Ebner, T.; Terasaki, T., Simultaneous absolute protein quantification of transporters, cytochromes P450, and UDP-glucuronosyltransferases as a novel approach for the characterization of individual human liver: comparison with mRNA levels and activities. Drug Metab Dispos 2012, 40, (1), 83–92.

6. Vogel, C.; Marcotte, E. M., Insights into the regulation of protein abundance from proteomic and transcriptomic analyses. Nature reviews. Genetics 2012, 13, (4), 227–32.

7. King, C.; Scott-Horton, T., Pyrosequencing: a simple method for accurate genotyping. J Vis Exp 2008, (11).

8. Pratt, J. M.; Simpson, D. M.; Doherty, M. K.; Rivers, J.; Gaskell, S. J.; Beynon, R. J., Multiplexed absolute quantification for proteomics using concatenated signature peptides encoded by QconCAT genes. Nat Protoc 2006, 1, (2), 1029–43.

9. Glatter, T.; Ludwig, C.; Ahrne, E.; Aebersold, R.; Heck, A. J.; Schmidt, A., Large-scale quantitative assessment of different in-solution protein digestion protocols reveals superior cleavage efficiency of tandem Lys-C/trypsin proteolysis over trypsin digestion. J Proteome Res 2012, 11, (11), 5145–56.

10. Shi, J.; Wang, X.; Lyu, L.; Jiang, H.; Zhu, H. J., Comparison of protein expression between human livers and the hepatic cell lines HepG2, Hep3B, and Huh7 using SWATH and MRM-HR proteomics: Focusing on drug-metabolizing enzymes. Drug Metab Pharmacokinet 2018, 33, (2), 133–140.

11. MacLean, B.; Tomazela, D. M.; Shulman, N.; Chambers, M.; Finney, G. L.; Frewen, B.; Kern, R.; Tabb, D. L.; Liebler, D. C.; MacCoss, M. J., Skyline: an open source document editor for creating and analyzing targeted proteomics experiments. Bioinformatics 2010, 26, (7), 966–8.

12. Vizcaino, J. A.; Csordas, A.; del-Toro, N.; Dianes, J. A.; Griss, J.; Lavidas, I.; Mayer, G.; Perez-Riverol, Y.; Reisinger, F.; Ternent, T.; Xu, Q. W.; Wang, R.; Hermjakob, H., 2016 update of the PRIDE database and its related tools. Nucleic acids research 2016, 44, (D1), D447–56.

13. Wang, D.; Johnson, A. D.; Papp, A. C.; Kroetz, D. L.; Sadee, W., Multidrug resistance polypeptide 1 (MDR1, ABCB1) variant 3435C>T affects mRNA stability. Pharmacogenet Genomics 2005, 15, (10), 693–704.

14. Penno, M. B.; Dvorchik, B. H.; Vesell, E. S., Genetic variation in rates of antipyrine metabolite formation: a study in uninduced twins. Proc Natl Acad Sci U S A 1981, 78, (8), 5193–6.

15. Ozdemir, V.; Kalow, W.; Tang, B. K.; Paterson, A. D.; Walker, S. E.; Endrenyi, L.; Kashuba, A. D., Evaluation of the genetic component of variability in CYP3A4 activity: a repeated drug administration method. Pharmacogenetics 2000, 10, (5), 373–88.

16. Eichler, E. E.; Flint, J.; Gibson, G.; Kong, A.; Leal, S. M.; Moore, J. H.; Nadeau, J. H., Missing heritability and strategies for finding the underlying causes of complex disease. Nature reviews. Genetics 2010, 11, (6), 446–450.

17. Khan, Z.; Ford, M. J.; Cusanovich, D. A.; Mitrano, A.; Pritchard, J. K.; Gilad, Y., Primate Transcript and Protein Expression Levels Evolve Under Compensatory Selection Pressures. Science 2013, 342, (6162), 1100–1104.

18. Cenik, C.; Cenik, E. S.; Byeon, G. W.; Grubert, F.; Candille, S. I.; Spacek, D.; Alsallakh, B.; Tilgner, H.; Araya, C. L.; Tang, H.; Ricci, E.; Snyder, M. P., Integrative analysis of RNA, translation, and protein levels reveals distinct regulatory variation across humans. Genome research 2015, 25, (11), 1610–21.

19. Chick, J. M.; Munger, S. C.; Simecek, P.; Huttlin, E. L.; Choi, K.; Gatti, D. M.; Raghupathy, N.; Svenson, K. L.; Churchill, G. A.; Gygi, S. P., Defining the consequences of genetic variation on a proteome-wide scale. Nature 2016, 534, (7608), 500–5.

20. Wu, Y.; Williams, E. G.; Dubuis, S.; Mottis, A.; Jovaisaite, V.; Houten, S. M.; Argmann, C. A.; Faridi, P.; Wolski, W.; Kutalik, Z.; Zamboni, N.; Auwerx, J.; Aebersold, R., Multilayered genetic and omics dissection of mitochondrial activity in a mouse reference population. Cell 2014, 158, (6), 1415–30.

21. Urquhart, B. L.; Tirona, R. G.; Kim, R. B., Nuclear receptors and the regulation of drug-metabolizing enzymes and drug transporters: implications for interindividual variability in response to drugs. J Clin Pharmacol 2007, 47, (5), 566–78.

22. Martelli, P. L.; Fariselli, P.; Savojardo, C.; Babbi, G.; Aggazio, F.; Casadio, R., Large scale analysis of protein stability in OMIM disease related human protein variants. BMC Genomics 2016, 17, (2), 397.

23. Khan, Z.; Bloom, J. S.; Amini, S.; Singh, M.; Perlman, D. H.; Caudy, A. A.; Kruglyak, L., Quantitative measurement of allele-specific protein expression in a diploid yeast hybrid by LC-MS. Molecular systems biology 2012, 8, 602–602.

24. Lee, H.; Qian, K.; von Toerne, C.; Hoerburger, L.; Claussnitzer, M.; Hoffmann, C.; Glunk, V.; Wahl, S.; Breier, M.; Eck, F.; Jafari, L.; Molnos, S.; Grallert, H.; Dahlman, I.; Arner, P.; Brunner, C.; Hauner, H.; Hauck, S. M.; Laumen, H., Allele-specific quantitative proteomics unravels molecular mechanisms modulated by cis-regulatory PPARG locus variation. Nucleic acids research 2017, 45, (6), 3266–3279.

25. Schubert, O. T.; Rost, H. L.; Collins, B. C.; Rosenberger, G.; Aebersold, R., Quantitative proteomics: challenges and opportunities in basic and applied research. Nat. Protocols 2017, 12, (7), 1289–1294.

26. Achour, B.; Russell, M. R.; Barber, J.; Rostami-Hodjegan, A., Simultaneous quantification of the abundance of several cytochrome P450 and uridine 5’-diphosphoglucuronosyltransferase enzymes in human liver microsomes using multiplexed targeted proteomics. Drug Metab Dispos 2014, 42, (4), 500–10.

27. Achour, B.; Dantonio, A.; Niosi, M.; Novak, J. J.; Fallon, J. K.; Barber, J.; Smith, P. C.; Rostami-Hodjegan, A.; Goosen, T. C., Quantitative Characterization of Major Hepatic UDP-Glucuronosyltransferase Enzymes in Human Liver Microsomes: Comparison of Two Proteomic Methods and Correlation with Catalytic Activity. Drug Metab Dispos 2017, 45, (10), 1102–1112.

28. Schechter, I.; Berger, A., On the size of the active site in proteases. I. Papain. Biochem Biophys Res Commun 1967, 27, (2), 157–62.

29. Giansanti, P.; Tsiatsiani, L.; Low, T. Y.; Heck, A. J., Six alternative proteases for mass spectrometry-based proteomics beyond trypsin. Nat Protoc 2016, 11, (5), 993–1006.

30. Cheung, C. S. F.; Anderson, K. W.; Wang, M.; Turko, I. V., Natural Flanking Sequences for Peptides Included in a Quantification Concatamer Internal Standard. Analytical Chemistry 2015, 87, (2), 1097–1102.

